# Alternative Promoters Drive Transcriptomic Reprogramming and Prognostic Stratification in TNBC

**DOI:** 10.1101/2025.03.19.643579

**Authors:** Simran Jit, Kirti Jain, Leepakshi Dhingra, Rahul Kumar, Sherry Bhalla

## Abstract

Transcriptional regulation frequently involves alternative promoters, yet the distinct regulatory mechanisms governing alternative versus reference promoters remain poorly understood in Triple-Negative Breast Cancer (TNBC), a high-risk breast tumor subtype. It is worth emphasizing that, despite the availability of extensive short-read sequencing data, the impact of alternative promoter usage on TNBC transcriptome dynamics and patient survival remains underexplored. The current study leverages RNA sequencing data from the publicly available TNBC tumor samples (360) and adjacent normal samples (88) to identify TNBC-specific and subtype-specific active alternative promoters (AAPs). To further validate these findings, we integrated H3K4me3 ChIP-seq data, confirming key promoter switching events. We found that *HDAC9, RPS14,* and *EPN1* exhibit tumor-specific AAP expression, while *AKAP9* and *SEC31A* show basal subtype-specific promoter activity in TNBC. Beyond their transcriptional impact, we also explored the prognostic significance of AAPs. The alternative promoters of *HUWE1* and *FTX* were recognized as independent survival predictors in TNBC. Notably, multivariate analysis demonstrated that prognostic AAPs remained significant in predicting relapse-free survival (RFS) even after adjusting for copy number alterations (CNA) and mRNA-based subtypes. Our AAP-based prognostic model achieved C-index of 0.73 in training and 0.72 in validation, with AUROC of 0.72 and integrated Brier score of 0.09 in validation dataset. Our findings indicate that AAP activity serves as a crucial prognostic marker beyond traditional clinical parameters, enhancing patient stratification and risk assessment in TNBC. Understanding promoter switching events may further unveil novel therapeutic targets, paving the way for precision oncology strategies in highly aggressive breast tumors.

## Background

Triple-negative breast cancer (TNBC) presents a major obstacle in oncology, owing to its aggressive behaviour and lack of effective targeted therapies (1,2). It is marked by the absence of estrogen receptor (ER), progesterone receptor (PR), as well as human epidermal growth factor receptor 2 (HER2), which limits treatment options and contributes to its poor prognosis (3). TNBCs demonstrate significant heterogeneity and have been further classified into 4 to 6 molecular subtypes determined by gene expression profiles (4,5). This complexity is driven by different regulatory mechanisms, including transcriptional regulation, epigenetic alterations, and post-transcriptional modifications (5). However, the contribution of alternative promoters (AP) in TNBC remains primarily underexplored due to the limited availability of genome-wide studies focusing on promoters. Techniques such as H3K4me3 histone modification profiling, an epigenetic marker of active promoters, or CAGE (cap analysis of gene expression), which identifies transcription start sites (TSSs), are extremely limited in pan-cancer studies (6,7).

Positioned upstream of transcription start site (TSS), promoters serve as crucial factors in the regulation of transcription. They determine not only when and how actively a gene is transcribed but also influence the specific isoform that will be expressed (8,9). According to recent studies, half of the protein-coding genes in mammals are controlled by alternative promoters which underpins their contribution to an important dimension of the complexity of the human transcriptome (10). By modulating transcriptional activity and isoform selection, promoters play a central role in gene expression diversity and its functional implications in different biological contexts (10,11). It is well established that promoters exhibit tissue-specific activity, thereby contributing to differential gene expression (12). This phenomenon has been supported by multiple studies showing that alternative promoters modulate the gene expression in a way highly dependent on cellular environment, developmental time as well as tissue type (13–15). They are key regulators of the isoform expression pre-transcriptionally. The aberrant expression of alternative promoter usage across various genes, has been observed in cancer. Various studies emphasize on the regulatory control exerted by alternative promoters, which can produce diverse transcripts and protein products, thereby influencing cancer progression (16–18).

Pan-cancer study by Demircioğlu et al. has shown that specific alternative promoters can be differentially regulated based on tumor type and different tumor subtypes, reinforcing the notion that they are integral players in cancer biology and may serve as promising biomarkers for prognosis. For breast tumors, the *STAU2* gene’s 3’-most promoter showed significantly higher activity in the basal subtype of breast tumors in comparison to the other subtypes. In contrast, the other two active *STAU2* promoters displayed similar or greater activity in non-basal subtypes, indicating that the choice of promoter is associated with the tumor’s molecular characteristics (19). In 2022, Yang YS et al. identified that the transcription of a TNBC-specific transcript was activated by a specific bromodomain containing protein 4 (*BRD4*) bound superenhancer. This oncogenic transcript, referred to as macrophage receptor with collagenous structure-tumor-specific transcript (MARCO-TST), was linked to poor survival in TNBC patients (20).

In the current study, we explored the role of alternative promoters in TNBC and their potential as prognostic indicators of patient survival. Through the analysis of RNA-seq data from the TNBC patients, we detected both active and active alternative promoters (AAPs) that are specific to TNBC and its molecular subtypes. To ensure the reliability of these findings, we further validated our findings using H3K4me3 ChIP-seq data. Notably, the expression of certain alternative promoters was significantly associated with reduced relapse-free survival, underscoring their potential as independent prognostic biomarkers. These results highlight the molecular heterogeneity of TNBC and emphasize on the critical importance of understanding alternative promoter usage to advance personalized therapeutic approaches and improve clinical outcomes for TNBC patients.

## Materials and Methods

### Data Collection

The raw paired-end RNA-seq bulk data from 448 samples, including 360 primary TNBC tissues and 88 adjacent normals from Fudan University Shanghai Cancer Center (FUSCC), was obtained from the Sequence Read Archive (SRA) database through the accession number SRP157974 (21). Additionally, three independent datasets were obtained from Gene Expression Omnibus (GEO) for validation purposes: GSE58135, GSE85158, and GSE240671. The external dataset 1, GSE58135, included paired-end RNA-seq data from 40 primary TNBC tissues and 21 adjacent normal tissues (22). It was used to validate TNBC-specific and TNBC subtype-specific active alternative promoters (AAPs). To analyze the trimethylation patterns of histone H3 at the lysine 4 (H3K4me3) in the identified AAPs, we utilized the ChIP-seq dataset GSE85158 which is referred to as the external dataset 2 throughout the paper (23). It comprised H3K4me3 data from cell lines representing normal breast as well as different molecular subtypes of breast tumor. We downloaded the hg38 aligned bigwig files from the ChIP-Atlas web tool (https://chip-atlas.org/) (24–26). Furthermore, the prognostic AAPs identified in this study were validated using the external dataset 3, GSE240671, comprising a total of 122 breast tumor samples, including 27 treatment-naive TNBC samples (27).

### RNA-seq pipeline and processing

For the bulk RNA-seq datasets (SRP157974, GSE58135, and GSE240671) utilized in this study, raw FASTQ files were downloaded using the SRA Toolkit (v2.10.9). Quality assessment of the reads was conducted with FastQC (v0.12.1) (28), and MultiQC (v0.4) (29) was employed to aggregate individual QC reports into a consolidated summary, facilitating improved visualization and assessment of data quality. Subsequently, Trimmomatic (v0.39) (30) was used to remove adapter sequences and trim bases with an average quality score below 15. The resulting high-quality reads were aligned to the human reference genome (Gencode v44; GRCh38) using STAR (v2.7.10b) (31). The resulting BAM files were then processed to generate raw counts by employing the featureCounts program (32) from the Subread package (v2.0.6) (33). Finally, transcript isoform quantification was performed using RSEM (v1.3.3) (34), resulting in generation of a Fragments Per Kilobase of transcript per Million mapped reads (FPKM) expression matrix.

### ChIP-seq data retrieval and analysis

The hg38 aligned bigwig files corresponding to normal (MCF10A; 76N-F2V) and TNBC (HCC1937; MB436) retrieved from the ChIP-Atlas web tool were employed for the validation of AAP specific H3K4me3 signals. Track plots were generated using R packages, including Gviz (v1.42.1) (35), GenomicRanges (v1.50.2) (36), rtracklayer (v1.58.0) (37), and GenomicAlignments (v1.34.1) (36), to facilitate the visualization of H3K4me3 enrichment patterns across defined genomic regions.

### Classification of tumor samples into PAM50 subtypes

The intrinsic subtype information from the FUSCC metadata was used to classify the TNBC samples into basal and non-basal tumor subtypes. For the external dataset 1, raw FASTQ files were obtained for 39 Non-TNBC samples and processed following the steps outlined in the section ‘RNA-seq Pipeline and Processing’. PAM50 subtyping was done using the R package genefu (v2.32.0) (38), and the corresponding subtypes for the 40 TNBC samples were determined in three rounds of molecular subtyping. To ensure consistency, approximately 75% of the Non-TNBC samples were included in each round of PAM50 subtyping. The TNBC samples were then categorized into basal and non-basal subtypes for the detection of basal specific AAPs.

### Classification of TNBC Samples

To determine the subtypes of TNBC samples from the FUSCC cohort and the external dataset 1, we utilized the TNBCtype tool (http://cbc.mc.vanderbilt.edu/tnbc) (4,39). The FPKM matrix generated from RSEM was used as an input for the analysis, with Ensembl IDs converted to gene names using the gprofiler2 R package (v0.2.3) (40). The matrix was log2 normalized (log2 FPKM + 1) and anonymized by masking the sample IDs prior to input into the TNBCtype tool. For samples predicted as ER+ by the tool, the ESR1 expression was adjusted by replacing it with the median ESR1 expression from the remaining samples. The modified matrix was then re-submitted to the TNBCtype tool and the obtained TNBC type 6 was subsequently mapped to TNBC type 4 as defined by Lehmann et al (5).

### Promoter Identification and Quantification of promoter activity

Promoters across the FUSCC cohort and the external dataset 1 were identified using the R package proActiv (v1.14.0) (19). Promoter activity was quantified based on the Gencode genome annotation (Gencode v44) and splice-junction files from STAR. To assess promoter activity, single-exon transcripts/promoters were excluded, and promoters with counts of NA or zero in both conditions of interest were removed. We also excluded internal promoters from our analysis because the first exon in case of internal promoter overlaps with the internal exons in other isoforms. As a result, the reads that map to such exons cannot conclusively be assigned to internal promoters as they may originate due to transcription of other isoforms that utilize those specific exons internally (19,41).

Wilcoxon rank-sum test was applied on the gene expression; absolute and relative promoter activity. We implemented the approach by Dong Y et al and identified differentially expressed genes (DEGs) and differentially regulated promoters (DRPs) (42). The criteria of p-value<0.05 and |Log2(fold change)|>1 was used to identify DEGs. Whereas, for DRP identification, p-value<0.05 and |Log2(fold change)|>1.2 was considered.

### Detection of Active Alternative Promoters (AAPs)

To identify the TNBC-specific AAPs, we first assessed the gene expression along with the absolute promoter activity. The genes with two or more promoters (multi-promoter genes) were retained for the downstream analysis. We initially investigated the non-significant genes (p-value>0.05) having both up and down significant (p-value<0.05) DRPs in terms of absolute promoter activity. In order to obtain more robust candidates, we further incorporated the relative promoter activity (p-value<0.05) in addition to the above-mentioned parameters and identified the TNBC-specific AAPs. For the detection of subtype-specific AAPs, the gene expression and both absolute and relative promoter activity were evaluated. Further, in order to define prognostic AAPs, additional filters included: 1) Genes with at least 2 promoters having >10 junction reads in at least 60% of tumor samples; 2) Relative promoter activity >0.25 in 10% of tumor samples; and 3) Non-zero absolute promoter activity in 10% of tumor samples.

### Dimensionality Reduction and Clustering

The R package FactoMineR (2.11) (43) was used for principal component analysis (PCA) and Rtsne (0.17) (44) was used for t-Distributed Stochastic Neighbor Embedding (t-SNE) analysis on gene expression and the absolute promoter activity matrix.

### Functional Enrichment Analysis

Pathways associated with the identified genes were analyzed using the R package gprofiler2 (v0.2.3). Annotated gene sets, including Hallmark, C2 curated (BioCarta, KEGG_Legacy, and Reactome), C5 Gene Ontology (GO), C6 oncogenic signature, and C7 immunologic signature, were fetched from the Molecular Signatures Database (MSigDB) (45). The differentially expressed genes (DEGs), differentially regulated promoter genes (DRPGs), and their overlapping genes were analyzed separately for upregulated and downregulated genes. These gene sets were then queried against the annotated pathways. For each gene set, the top 10 significant pathways were identified, and relevant pathways were selected for downstream visualization. Bubble plots for upregulated and downregulated genes were generated using the R package ggplot2 (v3.5.1) (46).

### Motif Analysis

To unmask the probable transcription factors that may act as upstream regulators of the promoter switching events, we performed the *de-novo* motif analysis via the Regulatory Sequence Analysis Tools (RSAT) web server (47,48). We considered sequences 200 bp upstream of the transcription start site (TSS) of promoters for the analysis. We compared the enrichment of motifs in the obtained AAP to other promoters of that gene in each case, to highlight the motifs that significantly contribute to these specific alternative promoters. The databases JASPAR (49), HOMER (50) and ENCODE (51) were employed for motif analysis. The following filters were applied to obtain robust and significant results: 1) Correlation>0.50; 2) e-value≤1.5×10^-4^; 3) Duplicate motifs removed by retaining the one with higher correlation.

### Survival Analysis

We employed the caret R package (v6.0-94) (52) to perform a stratified split of the dataset based on mRNA subtypes. In the training dataset, we dichotomized the activity of 120 promoters identified using the criteria described in the section ‘Detection of Active Alternative Promoters (AAPs)’ and the expression of their corresponding 56 genes into high and low categories. The maxstat R package (v0.7-25) (53) was employed for determining the optimal cutpoint for promoter activity and gene expression that maximized the difference in survival outcomes between the two groups.

To mitigate potential bias in defining cutoff values, we calculated the cutpoints across 5,000 random iterations, each using one-third of the samples, and derived the median cutoff value for each promoter and gene from these iterations. This process provided a maximally selected rank statistics cutoff value for promoter activity and gene expression.

Univariate survival analysis was performed using the Cox proportional hazards regression model from the survival R package (v3.2-13) (54,55) to assess relapse-free survival (RFS) in TNBC patients for each promoter and its corresponding gene. From the training dataset, those genes with AAPs were identified which showed statistically significant prognostic value for RFS (p-value<0.05), while the corresponding gene expression levels were not statistically significant (p-value>0.10).

We applied the cutoff values calculated from the training dataset to the validation dataset and repeated the evaluation process. Kaplan-Meier survival analysis was conducted using the survival R package, while multivariate survival analysis was done via the ggforestplot R package (v0.1.0). This analysis accounted for clinical parameters, including mRNA subtype and Copy number alterations (CNA) subtype, in combination with the validated prognostic AAPs.

### Construction of relapse-associated risk assessment model

To develop a relapse-associated risk assessment model in the FUSCC dataset, we analyzed various clinical features available in the metadata, including tumor size, Ki67 index, different copy number subtypes (Chr8p21 deletion, Chr9p23 amplification, Chr12p13 amplification, Chr13q34 amplification, Chr20q13 amplification, and Low-CIN), and age at surgery. Univariate survival analysis was performed for each selected feature to identify clinical parameters most strongly associated with patient survival.

We constructed multiple survival models, including LASSO, elastic net, and random forest, and evaluated their performance using metrics such as concordance index (c-index), integrated Brier score (IBS), and time-dependent area under the receiver operating characteristic (AUROC) curve. Model construction and evaluation were carried out using the scikit-survival (0.23.0) library in Python (3.12.2). To minimize sampling bias, all steps were repeated across ten different random states, and the mean performance metrics were reported as the final results.

## Results

### The Landscape of Active Promoter in TNBC

The promoter region plays a fundamental role in regulating the transcription. A large number of the genes in human genome are co-regulated by multiple, distinct promoters. Leveraging the approach outlined by Demircioğlu et al. (19) we identified the activity of 77,915 potential promoters within the FUSCC cohort of 360 TNBCs and 88 adjacent normal samples. Single-exon transcripts without quantifiable promoter activity were excluded, along with the internal promoters, resulting in the identification of promoter activities for 47,657 promoters across the samples, corresponding to 37,909 genes. Notably, 17.5% of these genes were regulated by two or more distinct promoters, while 82.5% were controlled by a single promoter (Fig 1A). Among these multi-promoter genes, 12.6% had two promoters, 3.2% had three promoters, and the remaining 1.7% were associated with four or more promoters (Supplementary Fig S1A). For the downstream analysis of alternative promoters in TNBC and its subtypes, our study specifically focused on these multi-promoter genes.

**Fig 1.**
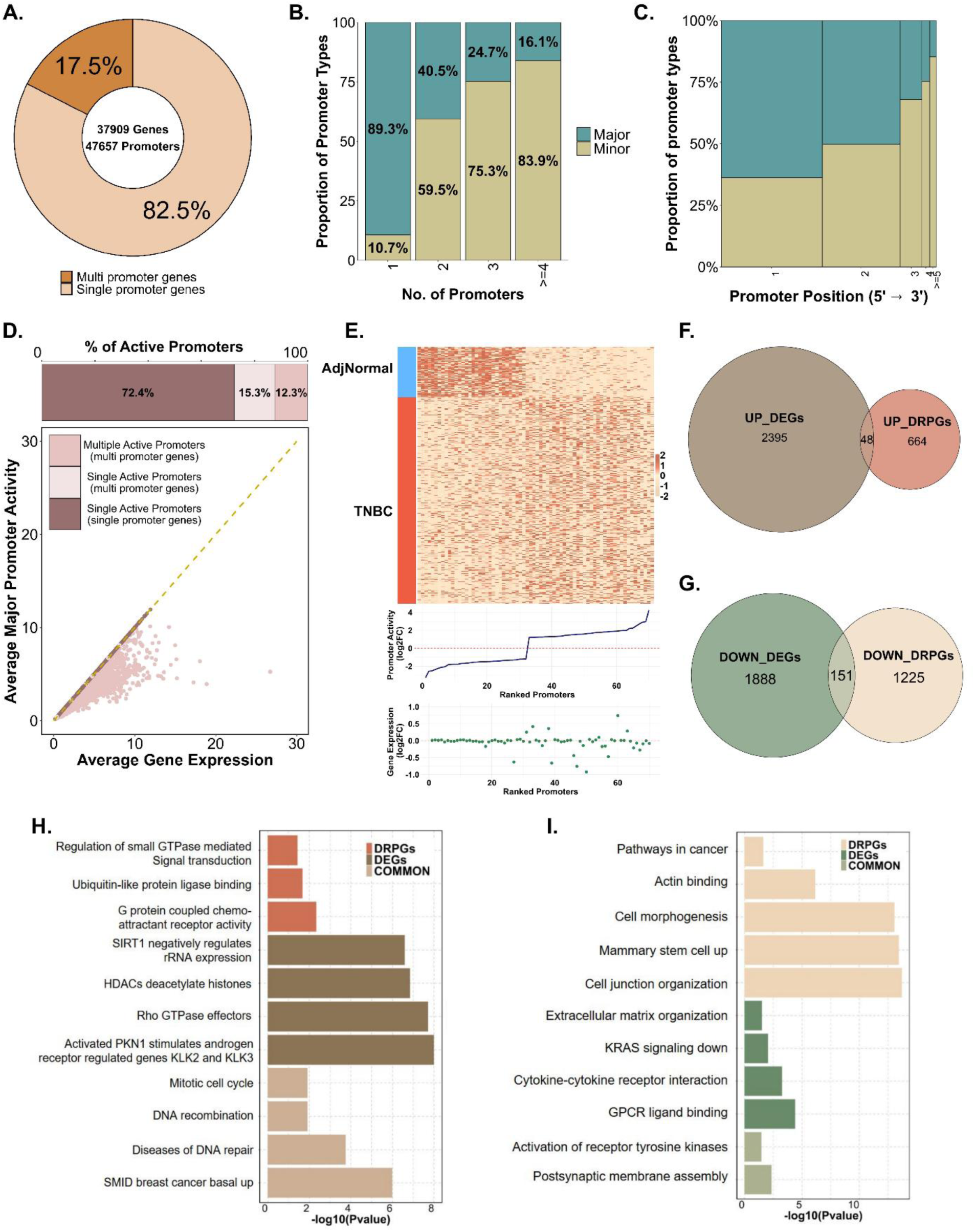
Estimation of Promoter activity using RNA-seq data. **A.** The total number of single and multi-promoter genes identified in FUSCC dataset. **B.** Proportions of major and minor promoters. **C.** The proportion of promoter types with respect to the position in gene. **D.** Number of single and multiple active promoters; and the comparison of average major promoter activity with average gene expression. Minor promoters of the genes uncover additional information. **E.** Heatmap showing the absolute promoter activity of DRPs in TNBC (n=360) and adjacent normal samples (n=88). The plots below highlight a distinctive pattern shown by promoter activity but not by the corresponding gene expression. **F.** Overlap between upregulated differentially expressed genes (DEGs) and the genes of upregulated differentially regulated promoters (DRPs). **G.** Common genes between downregulated DEGs and DRPGs. **H & I.** Bubble Plots representing enriched biological processes in upregulated and downregulated genes obtained.

The promoters were then classified into major and minor categories depending on the absolute promoter activity level (19). The promoter with the highest mean absolute activity across samples was defined as the major promoter of that gene. While, the promoters with average activity less than 0.25 were regarded as the inactive promoters and all the other remaining promoters of that gene constituted the minor promoters (Fig 1B). In multi-promoter genes, we observed that the proportion of minor promoters increased progressively with the number of promoters per gene. Typically, in the lack of regulatory genomics information, the first promoter of a gene is regarded as the active one. However, our findings reveal that the major promoter can be located at any position within a gene (Fig 1C). We found that the proportion of genes with promoters located downstream of first position decreases considerably, as shown by the narrowing of the bars in Fig 1C. Interestingly, 43.1% of major and 64% of minor promoters were positioned downstream of the first transcription start site (TSS) (Supplementary Fig S1B, C). This highlights the importance of RNA-seq data in offering dynamic functional insights and contextual information that static genome annotations alone cannot provide.

Certain promoters become active under specific conditions, and these promoters cannot be distinguished using traditional gene-level expression analysis. While the promoter activity typically aligns with overall gene expression, it reveals additional regulatory insights, particularly in cases where multiple promoters are active within a single gene. In our cohort, 12.3% of multi-promoter genes had multiple active promoters (Fig 1D). The significance of these promoters was reflected in comparison of the average major promoter activity with average gene expression. As shown in Figure 1D, the promoter activity for some promoters was found to be lower than the total gene expression, indicating that other promoters within the gene contribute more significantly to its expression, and that the observed promoter is less active in the given condition. This emphasizes the significance of promoter-level analysis in uncovering gene regulation within context-specific scenarios. It highlights that total gene expression is not solely driven by a single major promoter; often overlooked “minor promoters” also play a crucial role in contributing to overall gene activity.

Next, we screened for the differentially expressed genes (DEGs) and the differentially regulated promoters (DRPs) based on the gene expression and absolute promoter activity between TNBC and adjacent normal samples. In total, we identified 4,482 DEGs and 2,624 DRPs corresponding to 2032 unique genes. The heatmap in Fig 1E illustrates the ability of absolute promoter activity of DRPs to distinguish TNBC samples from adjacent normal tissues. Notably, the plots below the heatmap reveal a distinct separation at the promoter activity level, which becomes less pronounced when examining the gene expression of the same set of genes. This suggests that promoter-level variations are not reflected at the gene expression level and focusing solely on the gene expression may ultimately lead to the masking of critical insights. We next analyzed the overlap between upregulated DEGs and genes with upregulated DRPs (DRPGs), as well as downregulated DEGs and genes with downregulated DRPs (Fig 1F, G). This analysis revealed 48 overlapping genes in upregulated group and 151 in downregulated group. Functional enrichment analysis was conducted on these genes to identify the pathways they represent.

There was minimal overlap between the upregulated DEGs and genes with upregulated DRPs, as well as between the downregulated DEGs and genes with downregulated DRPs. The upregulated DRPGs were primarily enriched in pathways such as ubiquitin-like protein ligase binding and G-protein-coupled chemoattractant receptor (GPCR) signaling, while upregulated DEGs were predominantly associated with HDACs, SIRT1, and RHO GTPase activities. Interestingly, the overlapping genes were significantly enriched in pathways characteristic of the basal subtype of breast tumors, particularly those related to DNA repair (Fig 1H). For the downregulated DRPGs, pathways linked to cell junction organization and genes consistently upregulated in mammary stem cells were strongly enriched. Additional pathways related to cell morphogenesis and actin binding were also identified. Downregulated DEGs intersected with genes downregulated by KRAS activation, whereas the overlapping downregulated genes were primarily associated with receptor tyrosine kinase activation and postsynaptic membrane assembly (Fig 1I) The list of pathways obtained for the upregulated and downregulated genes is given in Supplementary Table S1, S2.

Overall, our results demonstrated that the promoter activity distinguishes TNBC samples from adjacent normal tissues, complementing gene expression analysis and offering deeper insights into tumorigenesis mechanisms that might be overlooked if we focus solely on the gene expression evaluation.

### Identification of TNBC-Specific Active Alternative Promoters

The comparative analysis of gene expression profiles between cancerous and normal tissues has played a pivotal role in identifying key cancer-associated genes and elucidating the biological pathways driving cancer progression (56). In this study, we aimed to determine if alternative promoters are specific to TNBC. Previous studies have indicated that alternative promoters could significantly influence transcriptional changes specific to cancer tissues (19,42). Such findings provide meaningful insights into the underlying disease mechanisms, providing the potential for more refined patient stratification and the development of improved treatment strategies. Guided by this understanding, we began to investigate the promoters that show an alternative usage between TNBC and adjacent normal samples in the FUSCC dataset. We concentrated on the multi-promoter genes having a non-significant change in expression between TNBC and adjacent normal samples. In order to detect promoter switching events, we fetched the genes having both upregulated and downregulated DRPs which showed significant difference in terms of their absolute promoter activity between TNBC and adjacent normal tissues. This approach led to the identification of 8 candidate genes (Fig 2A) (Supplementary Table S3). One of them, Histone Deacetylase 9 (*HDAC9*) gene, demonstrated a remarkable promoter switching wherein pr1077 was significantly downregulated and pr1079 was significantly upregulated in TNBC patients (Fig 2B, C). *HDAC9* gene encodes for the histone deacetylase 9 and has been known to promote tumor growth and progression (57). HDACs aid in the removal of acetyl group from the lysine residues of histones, thereby, playing a key role in regulating gene transcription and chromatin remodeling (58,59). According to recent studies, *HDAC* inhibitors have displayed potential to be employed for controlling the growth and progression of TNBC (58). We validated the lack of significant differential expression of *HDAC9* and the reduced promoter activity of pr1077 in TNBC samples in comparison to adjacent normal tissues using external dataset 1 (Fig 2D, E). We also validated our findings using an independent approach by analyzing the publicly available H3K4me3 ChIP-seq data, a widely recognized marker of active promoters. The trackplot presented in Fig 2F shows the pr1077 and pr1079 of *HDAC9* along with the mean bam file coverage and H3K4me3 peaks for these promoters in TNBC (HCC1937) and normal cell line (MCF10A). Analysis of H3K4me3 ChIP-seq data revealed consistently lower peak intensities for pr1077 and higher peak intensities for pr1079 in TNBC compared to adjacent normal tissues across all investigated tracks. In addition, the reduced activity of pr1077 in TNBC was also validated using another normal cell line (76N-F2V) wherein a similar pattern was recorded. This observation aligns with the observed differences in promoter activity, further supporting its differential regulation in TNBC.

**Fig 2.**
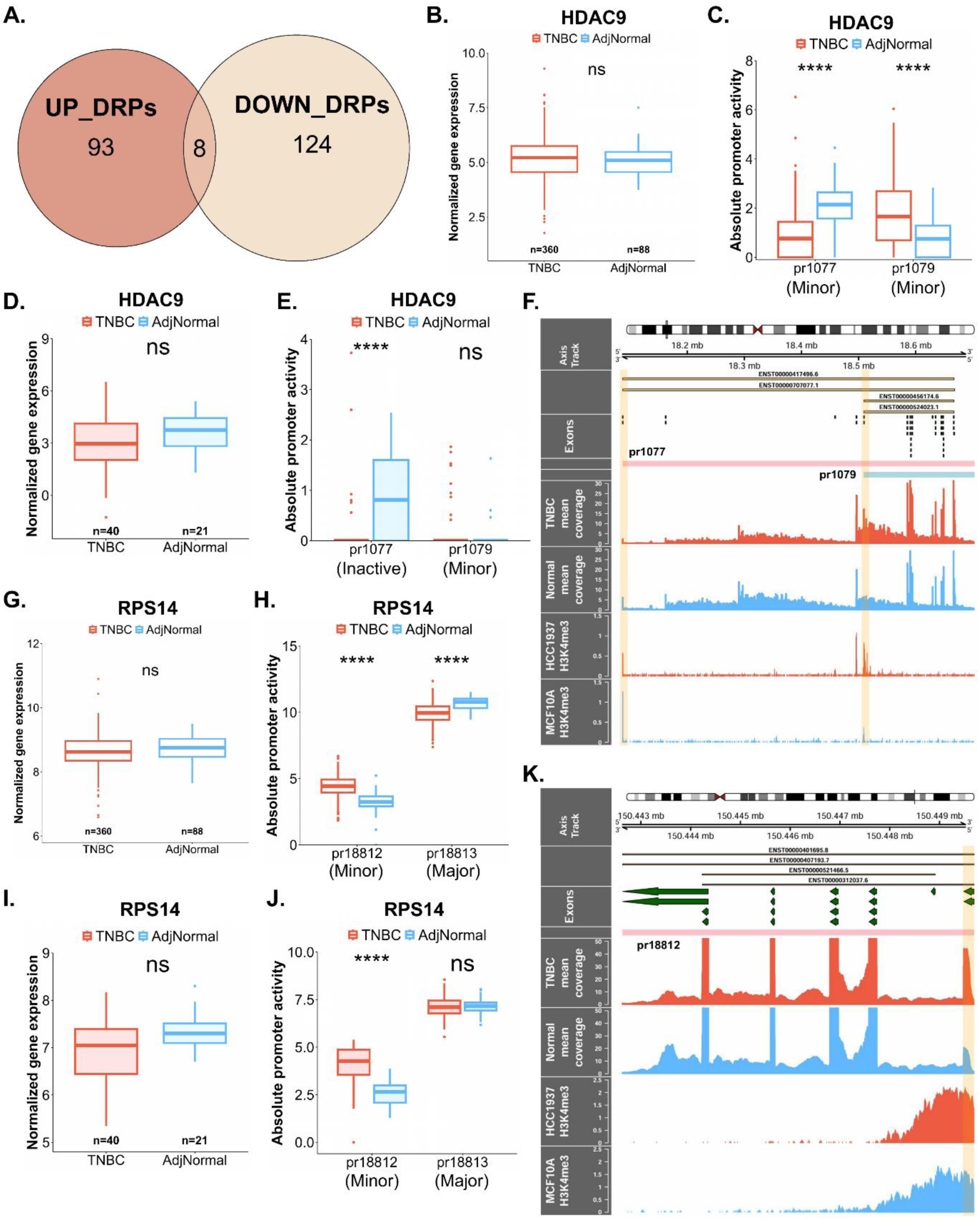
Identification of TNBC-specific AAPs. **A.** Venn diagram for the non-significant genes having significant up and downregulated DRPs as per absolute promoter activity in the FUSCC cohort. **B.** Non-significant gene expression of *HDAC9* between the TNBC and adjacent normals in FUSCC cohort. **C.** Boxplot showing significantly lower activity of *HDAC9* pr1077 and higher activity of pr1079 in TNBC patients in FUSCC cohort. **D & E.** Validation of *HDAC9* gene expression **(D)** and promoter activity **(E)** in the external dataset 1. **F.** Trackplot highlighting the promoter switching in *HDAC9* promoters in patients as well in the cell lines H3K4me3 ChIP-seq data. **G & H.** Status of *RPS14* gene expression **(G)** and absolute promoter activity **(H)** of its corresponding promoters in FUSCC. **I.** Validation of gene expression of *RPS14* in external dataset 1. **J.** pr18812 shows higher activity in TNBC in the external validation dataset 1. **K.** Trackplot showing the mean bam coverage and H3K4me3 ChIP-seq peaks for *RPS14* AAP in TNBC (HCC1937) and normal (MCF10A) cell lines.

In order to capture more robust TNBC-specific AAPs, we incorporated the relative promoter activity along with the previously applied criteria. Relative promoter activity is defined as the ratio of its absolute activity to the total absolute activity of all the promoters of that gene (19). We explored the genes with promoters exhibiting significant variation in both absolute and relative promoter activity, while showing no significant changes in their corresponding gene expression between TNBC and adjacent normal samples. Using this approach, we identified 217 candidate AAPs in the FUSCC dataset, of which 14 were validated in an external dataset 1. Principal Component Analysis (PCA) of the absolute promoter activity for these 14 AAPs revealed a clear distinction between TNBC and adjacent normal samples (Supplementary Fig S2A). In contrast, when PCA was performed using the gene expression of the captured genes, it failed to discriminate between TNBC and adjacent normal tissues, with all samples clustering together irrespective of their identity (Supplementary Fig S2B). In the FUSCC cohort, among the total promoters analyzed, 23.5% were classified as minor promoters, while 76.5% were the major promoters. Interestingly, on investigating the promoter types for the obtained 14 TNBC-specific AAPs, we found that 78.6% of these were the minor promoters (Supplementary Fig S2C). This pinpoints that alternative promoters are most commonly the minor promoters of a gene (19) and underscores their significance in providing meaningful insights in TNBC. Subsequently, we sought to examine the number of transcripts transcribed from the TNBC-specific AAPs and observed that 57.1% activated a single transcript, while the remaining activated more than one transcript (Supplementary Fig S2D).

Investigating the TNBC-specific AAPs closely, we found important candidate genes for which although the gene expression levels did not show a significant difference, the promoter activity of the AAP was elevated significantly in TNBC samples in comparison to adjacent normals. For instance, in the case of ribosomal protein S14 (*RPS14*) gene, the absolute promoter activity of pr18812 showed an upregulation specifically in TNBC while the gene expression failed to demonstrate a significant variation (Fig 2G, H). *RPS14* is a key structural protein of the 40S ribosomal subunit that contributes to the process of ribosomal biogenesis. Previous studies have shown *RPS14* to be an important stimulator of cell proliferation ultimately fueling tumor growth (60). It has been reported that *RPS14* is responsible for the human 5q-syndrome, a bone marrow disorder which is attributed to the interstitial deletion of chromosome 5 long arm (61). Zhou et al in 2013, demonstrated that *RPS14* plays a pivotal role in activating p53 activity via the inhibition of MDM2 by binding to its central domain (62). This binding interferes with the E3 ubiquitin ligase activity of MDM2 subsequently leading to cell-cycle arrest as well as inhibition of growth (63). Our findings on the *RPS14* gene and its corresponding minor promoter corroborated well in the external dataset 1 (Fig 2I, J). The higher activity of pr18812 was also validated through the H3K4me3 ChIP-seq data. The trackplot presented in Fig 2K shows the pr18812 of *RPS14* along with the mean bam file coverage and H3K4me3 peaks. Higher peak intensities for pr18812 were recorded in TNBC mean bam file coverage and TNBC cell line tracks (HCC1937). We also looked at all the three different promoters of *RPS14* and observed that pr18813, being the major promoter, showed a marked increase in mean bam coverage (Supplementary Fig S2E). Owing to this, we focussed on pr18812 individually to attain a better understanding of the peak signals in all tracks.

Apart from *HDAC9* and *RPS14*, we noted another interesting promoter switching event in the Epsin 1 gene (*EPN1*) which encodes a protein belonging to the epsin family and is a well-established endocytic adaptor (64). While gene expression levels showed no significant difference between TNBC and adjacent normal samples, the pr1475 of *EPN1* was significantly upregulated and pr1474 was significantly downregulated in TNBC samples relative to the adjacent normals (Supplementary Fig S2F, G). Investigation of *EPN1* gene expression and promoter activity in external dataset 1 revealed similar patterns with the minor promoter of the gene showing a significantly increased activity in TNBC patients (Supplementary Fig S2H, I). Our results were further supported by the H3K4me3 ChIP-seq analysis and higher peaks were detected at pr1475 in mean bam coverage and TNBC cell line (MB436) tracks (Supplementary Fig S2J).

After having detected meaningful promoter switching events, we aimed to gain a deeper insight into whether the observed events were driven by underlying changes in the upstream regulatory networks. To achieve this, we performed the *de-novo* motif analysis through the RSAT web server. We probed the enrichment of transcription factor motifs in AAPs as compared to the remaining active promoters of that gene. We observed certain significantly enriched motifs by querying the databases Homer, Encode and Jaspar for each of the TNBC-specific AAPs. We noticed important transcription factor motifs and the complete list of the detected motifs in AAPs of *HDAC9*, *RPS14* and *EPN1* is given in the Supplementary Table S4. Overall, deducing the important tumor-specific AAPs could enhance our understanding of the mechanisms driving tumor aggressiveness, allowing us to target them more efficiently and effectively.

### Identification of AAPs specific to subtypes

After identifying TNBC-specific AAPs, we sought to determine whether AAPs were also specific to different TNBC subtypes. This knowledge would enable a more in-depth exploration of promoter switching events in relation to the underlying heterogeneity of TNBC samples. Transcriptomic studies have delineated breast tumors into different intrinsic subtypes, each associated with unique clinical outcomes. For instance, the well known PAM50 classification, also referred to as the Prosigna Breast Cancer Prognostic Gene Signature Assay, divides the breast tumors into Luminal A, Luminal B, HER2-enriched, and Basal-like (65). Existing data suggests that a substantial majority of TNBCs, approximately 60–90%, fall into the basal breast tumor subtype, whereas only 11.5% of the Non-TNBCs coincide with basal tumors (66–68).

We were interested in determining basal specific AAPs and in order to achieve this, we analyzed the given basal and non-basal classification of TNBC samples in the metadata of the FUSCC dataset. There were 277 basal and 83 non-basal samples. The promoters in the two categories of interest were identified by running proActiv with this condition. We subsequently performed differential gene expression analysis and differential promoter activity analysis on absolute and relative promoter activity by applying wilcoxon rank-sum test. All internal promoters were excluded from the analysis and we retained only the multi-promoter genes. Non-significant genes (p-value>0.05) between basal and non-basal samples were fetched and their corresponding promoters which were significant in terms of both absolute and relative promoter activity (p-value<0.05) were identified as the basal subtype-specific AAPs. We obtained a total of 189 significant AAPs in the FUSCC dataset that fulfilled all the applied criteria.

Principal component analysis (PCA) was conducted on gene expression and absolute promoter activity of the candidate AAPs. We noticed that the gene expression of these AAPs did not differentiate the basal from non-basal samples, as they clustered together; however, absolute promoter activity showed slightly better separation of the samples (Supplementary Fig S3A, B).

The validation of the basal AAPs was carried out in the external dataset 1 wherein, PAM50 subtyping was performed on the 40 TNBC and 39 Non-TNBC samples using the genefu R package. We segregated the TNBCs into basal (n=35) and non-basal (n=5) categories and identified the promoters by running proActiv. DEGs and DRPs were identified by applying wilcoxon rank-sum test. A total of 114 AAPs were uncovered in basal vs non-basal samples in the external dataset 1 by applying the same filters as applied in the FUSCC dataset.

Among the captured basal specific AAPs, 6 were validated in the external dataset 1. PCA revealed that relative to gene expression, absolute promoter activity of the 6 basal AAPs provided a more effective separation of basals from non-basal samples (Supplementary Fig S3C, D). Figure 3A presents a heatmap of the absolute promoter activity of the 6 selected AAPs and it highlights the ability of these promoters to distinguish between the basal and non-basal samples. The first AAP belonged to the gene *AKAP9* and our analysis revealed that its gene expression demonstrated a non-significant difference whereas its promoter pr10005 was significantly downregulated in basal samples as compared to the non-basal samples both in the discovery cohort (FUSCC) (Fig 3B, C) as well as in the external dataset 1 (Fig 3D, E).

**Fig 3.**
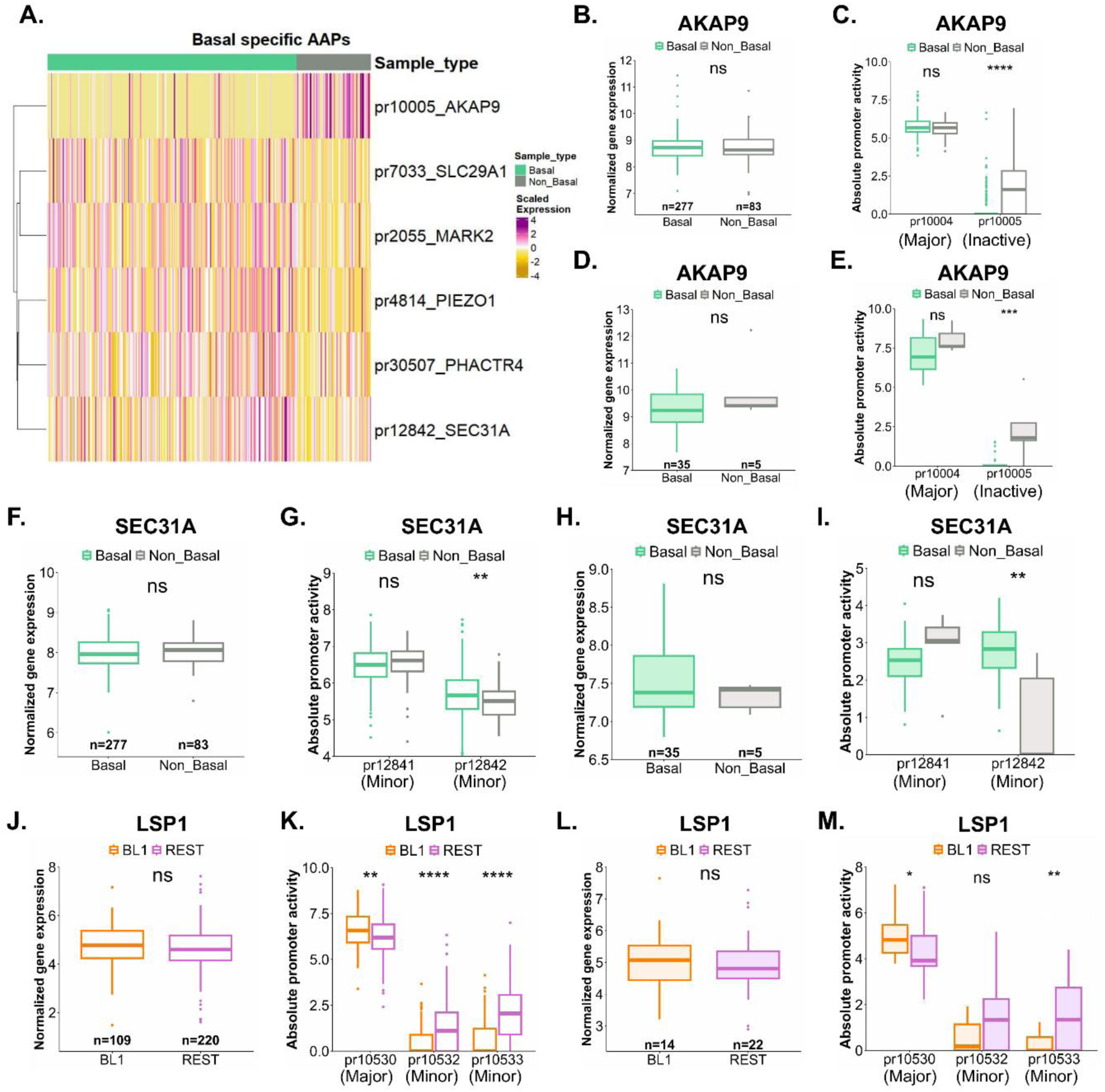
Identification of subtype-specific AAPs. **A.** Heatmap shows that the absolute promoter activity of the 6 basal specific AAPs notably segregates the basals from non-basal samples in the FUSCC cohort. **B.** Gene expression of *AKAP9* gene in FUSCC dataset. **C.** pr10005 shows a significant downregulation in basal samples in FUSCC. **D & E.** Validation of *AKAP9* gene expression **(D)** and absolute promoter activity **(E)** of its promoters in the external dataset 1. **F.** A non-significant difference in gene expression detected for the gene *SEC31A* in FUSCC. **G.** The activity of pr12842 shows significant upregulation in basal samples in the FUSCC dataset. **H & I.** Validation plots for *SEC31A* in the external dataset 1 for the gene expression **(H)** and absolute promoter activity **(I)**. **J.** Box plot depicting the variation in gene expression across BL1 and rest of the TNBC subtypes for *LSP1* gene in FUSCC. **K.** The minor promoters pr10532 and pr10533 show significantly low expression in the BL1 subtype in FUSCC. **L & M.** Gene expression **(L)** and promoter activity **(M)** for *LSP1* gene corroborate well in external dataset 1.

Another interesting candidate was highlighted in the *SEC31A* gene which encodes a component belonging to the coat protein complex II (COPII) and is essential in the process of cellular trafficking. We found that the gene expression of *SEC31A* did not change significantly (Fig 3F) but one of its promoter pr12842 was significantly high in the basal samples in contrast to non-basal samples (Fig 3G). We observed a similar trend in the external dataset 1 as well (Fig 3H, I).

Lehmann et al. (4,5) proposed that TNBCs display a higher degree of heterogeneity than previously recognized. In light of this, we further stratified the TNBCs into the 4 subtypes, namely BL1, BL2, LAR and M, to more effectively delineate their differences. We aimed to uncover potential promoter switching events specific to these subtypes. The distribution of patients in TNBC type 4 and PAM50 subtypes in FUSCC and external dataset 1 can be found in the Supplementary Table S5. A total of 214 BL1-specific AAPs were detected in the FUSCC dataset which belonged to 151 non-significant, unique genes. Parallelly, in the external dataset 1, we identified 80 AAPs corresponding to 74 unique genes. Out of all the obtained BL1 AAP candidates in the FUSCC dataset, 9 were successfully validated in the external dataset 1.

One of the interesting BL1-specific AAP was found in the gene Lymphocyte-specific protein 1 (*LSP1*). A non-significant variation in gene expression was detected between BL1 and the rest of the TNBC subtypes in the FUSCC cohort (Fig 3J). On comparing the absolute promoter activity of *LSP1* promoters, we found that pr10530 was significantly upregulated while the minor promoters pr10532 and pr10533 were significantly downregulated in the BL1 subtype of TNBC (Fig 3K). These patterns corroborated well in the external dataset 1 (Fig 3L, M). Further evaluation of all the TNBC subtypes separately for *LSP1* revealed that a non-significant gene expression difference existed between BL1 and LAR samples (Supplementary Fig S3E). We observed that the absolute promoter activity of both the minor promoters of *LSP1* was significantly lower in BL1 even when compared to all the different subtypes separately (Supplementary Fig S3F). We recorded a similar trend in external dataset 1 for the *LSP1* gene expression and promoter activity (Supplementary Fig S3G, H). *De-novo* motif analysis was performed for these subtype-specific alternative promoters and various differential transcription factor motifs were identified from Encode, Homer and Jaspar databases for each of the AAPs. The complete list of obtained motifs can be found in Supplementary Table S6.

### Prognostic Significance of Alternative Promoter Activity in TNBC

To investigate the prognostic significance of promoter activity in triple-negative breast cancer (TNBC), we analyzed relapse-free survival (RFS) data from the FUSCC cohort, which included 358 TNBC cases. Following the AAP selection criteria, we identified 120 active alternative promoters (AAPs) across 56 genes. The cohort was divided into training (70%) and validation (30%) sets, with each subset maintaining proportional representation of mRNA subtypes. Univariate survival analysis identified five overlapping genes, with *HUWE1* and *FTX* remaining significant after adjustment for clinical factors (Supplementary table S8, S9).

Notably, we found that reduced promoter activity of *HUWE1* (Fig 4A-C) and increased promoter activity of *FTX* (Fig 4D-F; Supplementary Fig S4A) were both significantly associated with shorter RFS, despite the absence of similar associations at the gene expression level. *HUWE1*, an E3 ligase playing a role in DNA repair, autophagy, and apoptosis, has been identified to function both as an oncogene and tumor suppressor across cancers (69,70). Kaplan-Meier analysis revealed that lower promoter activity of *HUWE1* (pr3004) was linked to higher relapse rates (log-rank p-value<0.0001), even though there was no correlation with HUWE1 gene expression.

**Fig 4.**
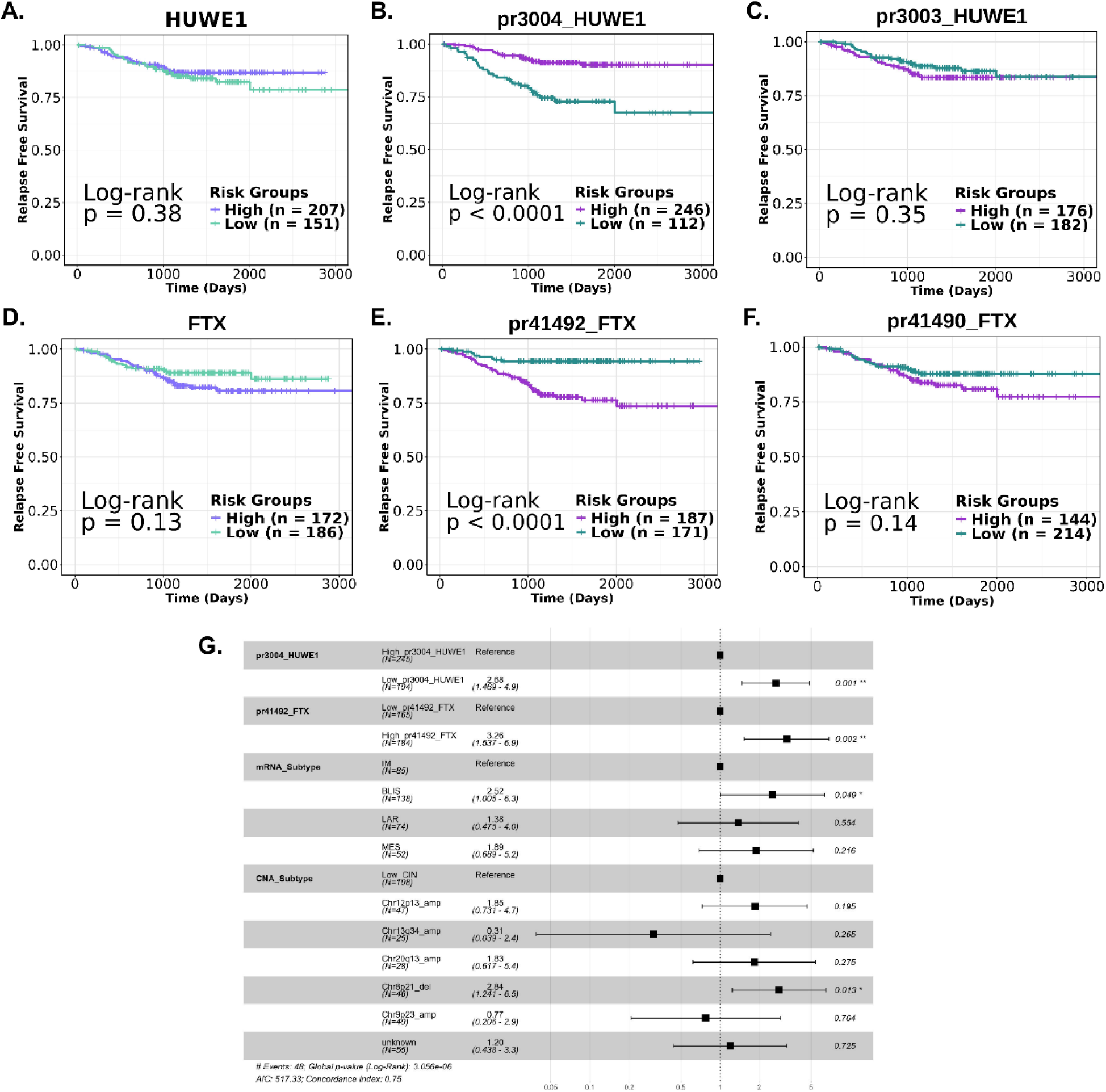
Promoter activity enhances prognostic stratification for patients with TNBC. **A.** RFS stratification by HUWE1 gene expression. **B,C.** RFS stratification by HUWE1 promoter activity, with significant stratification observed for its promoter pr3004. **D.** RFS based on FTX gene expression, **E,F.** RFS based on FTX promoter activity, with significant stratification for its promoter pr41492. **G.** Multivariate survival analysis demonstrating the impact of FTX and HUWE1 promoter activity, adjusted for other clinical subtypes.

Similarly, *FTX*, a non-coding RNA involved in epigenetic regulation and cellular metabolism, exhibited elevated promoter activity (pr41492) associated with increased relapse rates (log-rank p-value<0.0001), despite a lack of correlation of gene expression for RFS. Previous studies have pointed towards the role of *FTX* in tumor development and progression, although its utility as a therapeutic target remains unexplored (71).

Validation in the GSE240671 cohort (27 TNBC samples, high-vs low-risk residual cancer burden categories) revealed elevated *FTX* promoter (pr41492) expression in high-risk samples, albeit non-significant (Wilcoxon p-value=0.19) (Supplementary Fig S4B-E), possibly due to sample size limitations. *HUWE1* promoter (pr3004) activity remained constant across risk groups. Extending this analysis to a broader breast cancer cohort (n=95) further supported these trends, with high-risk patients showing elevated *FTX* promoter activity and low-risk patients showing increased *HUWE1* promoter activity, though differences were statistically non-significant, they align with our initial findings, thereby providing further support for these observations (Supplementary Fig S4F-I).

Through multivariate Cox regression analysis, we observed that both the identified prognostic promoters retained their significance for relapse-free survival (RFS), even after adjusting for confounding factors such as expression-based and copy number-based molecular subtypes. This finding highlights the independent prognostic value of promoter activity in TNBC (Fig 4G).

*De-novo* motif analysis comparing the pr3004 and pr3003 promoters of *HUWE1* uncovered several enriched motifs, including binding sites for key transcription factors such as ZBTB17 and ZNF519. According to the current evidence, high ZBTB17 expression has been found to be related to good survival in case of neuroblastoma patients (p-value=0.0048) (72). Furthermore, according to the research by Fan et al. in 2022, ZNF519 was identified as a key hub gene linked to long-term survival in the METABRIC cohort of breast cancer patients (73). Similarly, for *FTX* pr41492 one of the motifs obtained corresponded to the E74-like transcription factor (Elf5) which has been regarded as a tumour suppressive transcription factor (74). It is primarily expressed in the epithelial cells and serves as a key inhibitor of EMT. The list of transcription factor motifs obtained for prognostic AAPs is provided in the Supplementary Table S7.

### Alternative Promoter Activity as a survival risk biomarker in TNBC

To assess the clinical relevance of our selected prognostic promoters, pr41492 of *FTX* and pr3004 of *HUWE1*, we constructed three risk assessment models. The first model incorporated conventional clinical biomarkers, specifically tumor size and Ki-67 index score (Fig 5A); the second model utilized the selected prognostic promoter activities (Fig 5B); and the third model combined both sets of features (Fig 5C). Only tumor size and the Ki-67 index score were selected for the clinical model, as these were the only features that reached significance in univariate analysis and are routinely used in clinical settings to assess TNBC prognosis-related risk (Supplementary Table S10). Patients were classified into risk groups based on event frequency, and Kaplan-Meier analysis was used to evaluate model performance.

**Fig 5.**
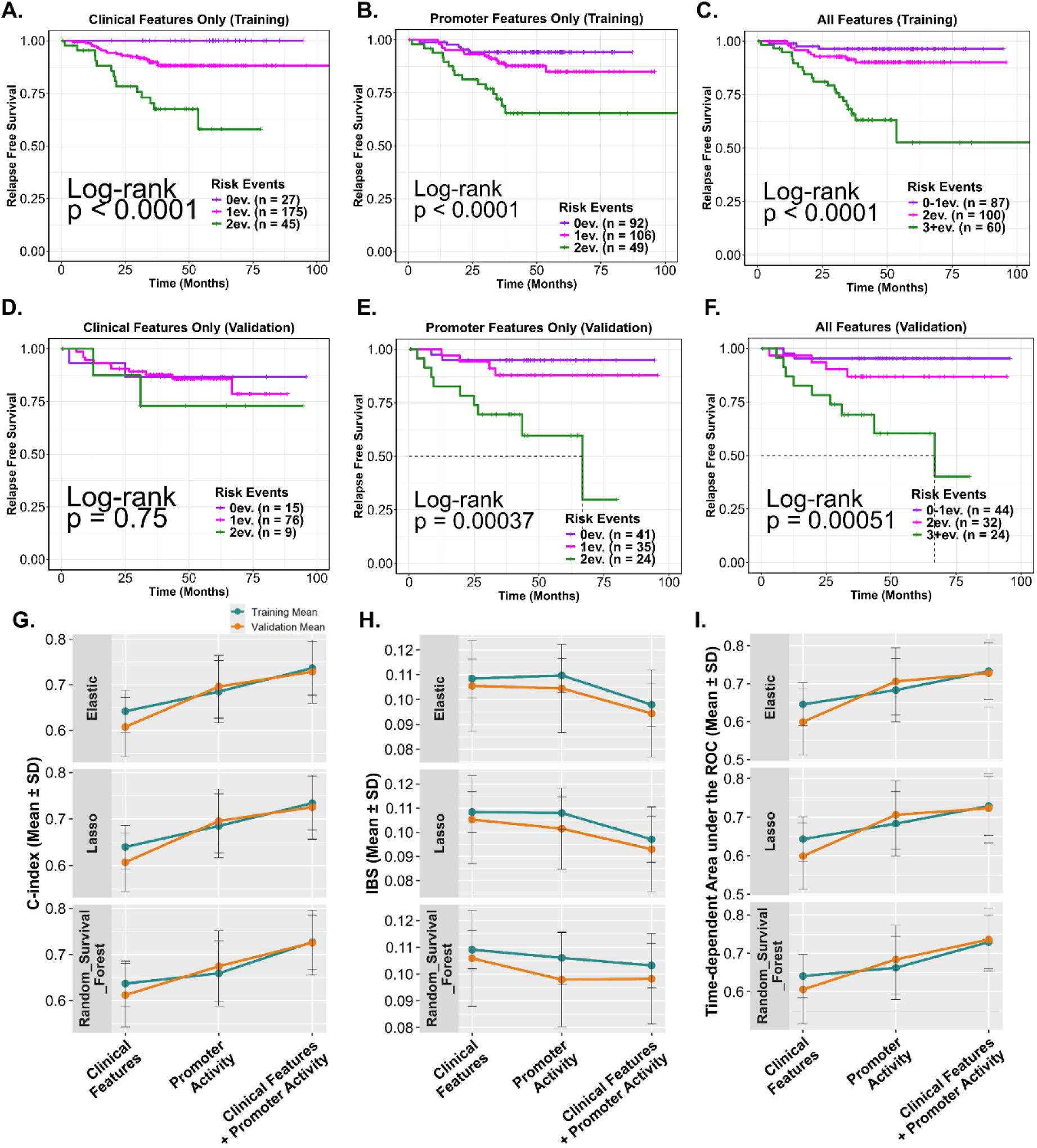
Evaluation of risk assessment models. **A.** Kaplan-Meier curves with dichotomized events of RFS in the training dataset using clinical features only **B.** With prognostic promoters activity only. **C.** Using the combination of clinical and prognostic promoter activity as features. **D, E, F.** RFS curves for the validation dataset. **G.** Evaluation of the model performance using Lasso, Elastic-Net, and Random Survival Forest (RSF) approaches, with the concordance index (C-index) as the metric. **H.** Assessment with Integrated Brier Score (IBS) as a measure. **I.** Using Time-dependent area under ROC as a metric for evaluation.

Results revealed that the first model, using only clinical biomarkers, did not significantly separate high-risk groups in the validation cohort (log-rank p-value=0.75) (Fig 5D). In contrast, the second model, which used promoter activity, significantly stratified high-risk patients in both training (log-rank p-value<0.0001) (Fig 5B) and validation datasets (log-rank p-value<0.00037) (Fig 5E). The third, combined model further improved patient stratification, achieving significant separation in both the training (log-rank p-value<0.0001) and validation datasets (log-rank p-value<0.00051) (Fig 5C, F; Supplementary Fig S5A, B).

We confirmed that the distribution of clinical features between training and validation cohorts was consistent, ruling out bias. Model performance was evaluated with concordance index (C-index), integrated Brier score (IBS), and time-dependent AUROC. Across all metrics, combining clinical and promoter activity features improved the models, with promoter activity alone outperforming clinical features when analyzed independently (Fig 5G-I). In summary, promoter activity-derived expression significantly enhances survival stratification over traditional clinical biomarkers in TNBC, underlining its value in refining risk assessment and informing patient management strategies.

## Discussion

The promoter region, a key DNA sequence where transcription factors bind, plays a vital role in regulating the gene expression and initiating transcription. While promoter studies have traditionally been conducted on smaller cohorts, the availability of RNA-seq data from a larger patient population offers a valuable opportunity to explore alternative promoters. In this study, we applied the method developed by Demircioğlu et al. (19) integrating RNA-seq data with genome annotation to define promoters in a larger cohort. Our analysis focuses on the complex role of alternative promoters (AP) in Triple-Negative Breast Cancer (TNBC), a subtype recognized for its highly aggressive nature and constrained therapeutic options. Our methodology utilized publicly available raw RNA-seq data from a large FUSCC cohort, complemented by ChIP-seq validation to confirm promoter activity, thereby offering a more comprehensive insight into the regulatory landscape of TNBC.

The identification of TNBC-specific and subtype-specific active alternative promoters (AAPs) reveals a complex regulatory landscape that may contribute to tumor heterogeneity and progression. Notably, genes like *HDAC9*, *RPS14* and *EPN1* displayed tumor-specific alternative promoter expression, whereas specific AAPs of *SEC31A* and *AKAP9* were associated with the basal and non-basal subtypes of TNBC, respectively. Further delving into detailed subtypes we also found that the minor promoters of *LSP1* gene were specifically associated with BL1 samples. This suggests that alternative promoter usage could serve as a mechanism for adapting to the tumor microenvironment, potentially influencing therapeutic responses.

Our study also highlights the prognostic significance of active alternative promoter (AAP) activity in TNBC. Notably, the promoter activity of *HUWE1* and *FTX*, two genes identified through our analysis, was significantly linked to shorter RFS, despite no changes in the corresponding gene expression. Reduced promoter activity of *HUWE1* promoter and an elevated activity of *FTX* promoter were associated with increased relapse rates, underlining the independent prognostic value of promoter activity in TNBC. Multivariate analysis confirmed that both *HUWE1* and *FTX* retained their significance even after adjusting for other clinical and molecular factors. Furthermore, when integrated into risk assessment models, promoter activity significantly improved survival stratification, outperforming conventional clinical biomarkers such as tumor size and Ki-67 index. This emphasizes the potential of AAPs in refining prognostic models and enhancing patient management strategies in TNBC. Our findings suggest that alternative promoter activity could serve as potential biomarkers for identifying high-risk patients in TNBC, offering valuable insights into the molecular mechanisms driving tumor progression.

Understanding the mechanisms behind promoter switching events opens avenues for targeted therapeutic strategies aimed at these regulatory elements. Given the potential impact of AAPs on tumor progression and patient prognosis, future research should be directed towards the elucidation of molecular pathways governing their activity. Additionally, exploring the therapeutic implications of targeting specific AAPs could lead to novel treatment strategies for TNBC patients. However, our study has few limitations including, uncertainty regarding the underlying mechanisms that stimulate or inhibit the activity of a promoter in TNBC samples. Detailed investigation about the influence of transcription factors on promoter activity would help uncover crucial regulatory mechanisms in the context of AAPs. Apart from this, experimental validation is necessary to complement the *in-silico* results.

## Conclusions

This study explores the role of alternative promoter (AP) activity in triple-negative breast cancer (TNBC) through the RNA-seq data from TNBC and adjacent normal samples. The study identifies TNBC-specific active alternative promoters (AAPs), validated using H3K4me3 ChIP-seq data. *HDAC9*, *RPS14*, and *EPN1* show tumor-specific AAP expression. We also identified several promoters showing basal subtype-specific activity. Notably, *HUWE1* and *FTX* AAPs are identified as independent predictors of prognosis. The findings suggest AP is a valuable prognostic marker beyond clinical parameters, enhancing patient stratification and risk assessment. This research highlights TNBC’s molecular heterogeneity and the importance of understanding AP for personalized therapeutic approaches.

## List of Abbreviations

TNBC: Triple-negative breast cancer
AAP: Active alternative promoter
AdjNormal: Adjacent normal
RNA-seq: RNA sequencing
ChIP-seq: Chromatin immunoprecipitation followed by sequencing
RFS: Relapse-free survival
C-index: Concordance index
IBS: Integrated Brier score
AUROC: Area under the receiver operating characteristic (ROC) curve
ER: Estrogen receptor
PR: Progesterone receptor
HER2: Human epidermal growth factor receptor 2
CAGE: Cap analysis of gene expression
TSS: Transcription start site
SRA: Sequence Read Archive
GEO: Gene Expression Omnibus
DEGs: Differentially expressed genes
DRPs: Differentially regulated promoters
PCA: Principal component analysis
t-SNE: t-Distributed Stochastic Neighbor Embedding
CNA: Copy number alteration
CIN: Chromosomal instability
BL1: Basal-like 1
BL2: Basal-like 2
LAR: Luminal androgen receptor
M: Mesenchymal
LASSO: Least Absolute Shrinkage and Selection Operator

## Declarations

### Ethics approval and consent to participate

Not applicable

### Consent for publication

Not applicable

### Availability of data and materials

The datasets analysed in the current study are available in the SRA repository under the accession number SRP157974 and in GEO database under the accession numbers GSE58135, GSE85158, and GSE240671.

### Authors Contribution

SB, SJ and KJ conceptualized the work. SB and SJ acquired the data and performed preprocessing of the raw data. SJ, KJ, SB analyzed the data. SJ, KJ, LD, RK and SB interpreted the data. The manuscript was written by SJ, KJ, LD and SB. All authors contributed to its editing and reviewed the final version.

## Acknowledgements

SJ acknowledges University Grants Commission (UGC) for fellowship support. KJ acknowledges CSIR for fellowship support. LD acknowledges CSIR-IGIB for support. SB acknowledges DBT/Wellcome Trust India Alliance Early Career Fellowship (ECF). We acknowledge the support from CSIR-IGIB for infrastructure and its IT department for computing facilities.

## References

1. Almansour NM. Triple-Negative Breast Cancer: A Brief Review About Epidemiology, Risk Factors, Signaling Pathways, Treatment and Role of Artificial Intelligence. Front Mol Biosci. 2022 Jan 25;9:836417.

2. Anders CK, Abramson V, Tan T, Dent R. The Evolution of Triple-Negative Breast Cancer: From Biology to Novel Therapeutics. Am Soc Clin Oncol Educ Book Am Soc Clin Oncol Annu Meet. 2016;35:34–42.

3. McGuire A, Lowery AJ, Kell MR, Kerin MJ, Sweeney KJ. Locoregional Recurrence Following Breast Cancer Surgery in the Trastuzumab Era: A Systematic Review by Subtype. Ann Surg Oncol. 2017 Oct 1;24(11):3124–32.

4. Lehmann BD, Bauer JA, Chen X, Sanders ME, Chakravarthy AB, Shyr Y, et al. Identification of human triple-negative breast cancer subtypes and preclinical models for selection of targeted therapies. J Clin Invest. 2011 Jul 1;121(7):2750–67.

5. Lehmann BD, Jovanović B, Chen X, Estrada MV, Johnson KN, Shyr Y, et al. Refinement of Triple-Negative Breast Cancer Molecular Subtypes: Implications for Neoadjuvant Chemotherapy Selection. PLoS ONE. 2016 Jun 16;11(6):e0157368.

6. Bernstein BE, Humphrey EL, Erlich RL, Schneider R, Bouman P, Liu JS, et al. Methylation of histone H3 Lys 4 in coding regions of active genes. Proc Natl Acad Sci U S A. 2002 Jun 25;99(13):8695–700.

7. Kodzius R, Kojima M, Nishiyori H, Nakamura M, Fukuda S, Tagami M, et al. CAGE: cap analysis of gene expression. Nat Methods. 2006 Mar;3(3):211–22.

8. Ayoubi TA, Van De Ven WJ. Regulation of gene expression by alternative promoters. FASEB J Off Publ Fed Am Soc Exp Biol. 1996 Mar;10(4):453–60.

9. Carninci P, Sandelin A, Lenhard B, Katayama S, Shimokawa K, Ponjavic J, et al. Genome-wide analysis of mammalian promoter architecture and evolution. Nat Genet. 2006 Jun;38(6):626–35.

10. Davuluri RV, Suzuki Y, Sugano S, Plass C, Huang THM. The functional consequences of alternative promoter use in mammalian genomes. Trends Genet. 2008 Apr 1;24(4):167–77.

11. Pankratova EV. Alternative promoters and the complexity of the mammalian transcritome. Mol Biol (Mosk). 2008;42(3):422–33.

12. Kamat A, Hinshelwood MM, Murry BA, Mendelson CR. Mechanisms in tissue-specific regulation of estrogen biosynthesis in humans. Trends Endocrinol Metab. 2002 Apr 1;13(3):122–8.

13. Hamaya Y, Suzuki A, Suzuki Y, Tsuchihara K, Yamashita R. Classification and characterization of alternative promoters in 26 lung adenocarcinoma cell lines. Jpn J Clin Oncol. 2022 Dec 3;53(2):97–104.

14. Kimura K, Wakamatsu A, Suzuki Y, Ota T, Nishikawa T, Yamashita R, et al. Diversification of transcriptional modulation: Large-scale identification and characterization of putative alternative promoters of human genes. Genome Res. 2006 Jan;16(1):55–65.

15. Maunakea AK, Nagarajan RP, Bilenky M, Ballinger TJ, D’Souza C, Fouse SD, et al. Conserved Role of Intragenic DNA Methylation in Regulating Alternative Promoters. Nature. 2010 Jul 8;466(7303):253–7.

16. Nepal C, Andersen JB. Alternative promoters in CpG depleted regions are prevalently associated with epigenetic misregulation of liver cancer transcriptomes. Nat Commun. 2023 May 11;14:2712.

17. Singer GA, Wu J, Yan P, Plass C, Huang TH, Davuluri RV. Genome-wide analysis of alternative promoters of human genes using a custom promoter tiling array. BMC Genomics. 2008 Jul 25;9:349.

18. Valcárcel LV, Amundarain A, Kulis M, Charalampopoulou S, Melnick A, San Miguel J, et al. Gene expression derived from alternative promoters improves prognostic stratification in multiple myeloma. Leukemia. 2021;35(10):3012–6.

19. Demircioğlu D, Cukuroglu E, Kindermans M, Nandi T, Calabrese C, Fonseca NA, et al. A Pan-cancer Transcriptome Analysis Reveals Pervasive Regulation through Alternative Promoters. Cell. 2019 Sep 5;178(6):1465–1477.e17.

20. Yang YS, Jin X, Li Q, Chen YY, Chen F, Zhang H, et al. Superenhancer drives a tumor-specific splicing variant of MARCO to promote triple-negative breast cancer progression. Proc Natl Acad Sci U S A. 2022 Nov 15;119(46):e2207201119.

21. Jiang YZ, Ma D, Suo C, Shi J, Xue M, Hu X, et al. Genomic and Transcriptomic Landscape of Triple-Negative Breast Cancers: Subtypes and Treatment Strategies. Cancer Cell. 2019 Mar 18;35(3):428–440.e5.

22. Varley KE, Gertz J, Roberts BS, Davis NS, Bowling KM, Kirby MK, et al. Recurrent read-through fusion transcripts in breast cancer. Breast Cancer Res Treat. 2014;146(2):287– 97.

23. Franco HL, Nagari A, Malladi VS, Li W, Xi Y, Richardson D, et al. Enhancer transcription reveals subtype-specific gene expression programs controlling breast cancer pathogenesis. Genome Res. 2018 Feb;28(2):159–70.

24. Oki S, Ohta T, Shioi G, Hatanaka H, Ogasawara O, Okuda Y, et al. ChIP-Atlas: a data-mining suite powered by full integration of public ChIP-seq data. EMBO Rep. 2018 Dec;19(12):e46255.

25. Zou Z, Ohta T, Miura F, Oki S. ChIP-Atlas 2021 update: a data-mining suite for exploring epigenomic landscapes by fully integrating ChIP-seq, ATAC-seq and Bisulfite-seq data. Nucleic Acids Res. 2022 Jul 5;50(W1):W175–82.

26. Zou Z, Ohta T, Oki S. ChIP-Atlas 3.0: a data-mining suite to explore chromosome architecture together with large-scale regulome data. Nucleic Acids Res. 2024 Jul 5;52(W1):W45–53.

27. Derouane F, Desgres M, Moroni C, Ambroise J, Berlière M, Van Bockstal MR, et al. Metabolic adaptation towards glycolysis supports resistance to neoadjuvant chemotherapy in early triple negative breast cancers. Breast Cancer Res BCR. 2024;26:29.

28. Andrews S. A quality control tool for high throughput sequence data. 2010; Available from: http://www.bioinformatics.babraham.ac.uk/projects/fastqc/

29. Ewels P, Magnusson M, Lundin S, Käller M. MultiQC: summarize analysis results for multiple tools and samples in a single report. Bioinformatics. 2016 Oct 1;32(19):3047– 8.

30. Bolger AM, Lohse M, Usadel B. Trimmomatic: a flexible trimmer for Illumina sequence data. Bioinformatics. 2014 Aug 1;30(15):2114–20.

31. Dobin A, Davis CA, Schlesinger F, Drenkow J, Zaleski C, Jha S, et al. STAR: ultrafast universal RNA-seq aligner. Bioinformatics. 2013 Jan;29(1):15–21.

32. Liao Y, Smyth GK, Shi W. featureCounts: an efficient general purpose program for assigning sequence reads to genomic features. Bioinformatics. 2014 Apr 1;30(7):923– 30.

33. Liao Y, Smyth GK, Shi W. The Subread aligner: fast, accurate and scalable read mapping by seed-and-vote. Nucleic Acids Res. 2013 May;41(10):e108.

34. Li B, Dewey CN. RSEM: accurate transcript quantification from RNA-Seq data with or without a reference genome. BMC Bioinformatics. 2011 Aug 4;12(1):323.

35. Hahne F, Ivanek R. Visualizing Genomic Data Using Gviz and Bioconductor. In: Mathé E, Davis S, editors. Statistical Genomics: Methods and Protocols. New York, NY: Springer; 2016. p. 335–51. Available from: 10.1007/978-1-4939-3578-9_16

36. Lawrence M, Huber W, Pagès H, Aboyoun P, Carlson M, Gentleman R, et al. Software for Computing and Annotating Genomic Ranges. PLOS Comput Biol. 2013 Aug 8;9(8):e1003118.

37. Lawrence M, Gentleman R, Carey V. rtracklayer: an R package for interfacing with genome browsers. Bioinformatics. 2009 Jul 15;25(14):1841–2.

38. Gendoo DMA, Ratanasirigulchai N, Schröder MS, Paré L, Parker JS, Prat A, et al. Genefu: an R/Bioconductor package for computation of gene expression-based signatures in breast cancer. Bioinformatics. 2016 Apr 1;32(7):1097–9.

39. Chen X, Li J, Gray WH, Lehmann BD, Bauer JA, Shyr Y, et al. TNBCtype: A Subtyping Tool for Triple-Negative Breast Cancer. Cancer Inform. 2012 Jan 1;11:CIN.S9983.

40. Kolberg L, Raudvere U, Kuzmin I, Vilo J, Peterson H. gprofiler2 -- an R package for gene list functional enrichment analysis and namespace conversion toolset g:Profiler. F1000Research. 2020 Nov 17;9:ELIXIR-709.

41. Zhang M, Sjöström M, Cui X, Foye A, Farh K, Shrestha R, et al. Integrative analysis of ultra-deep RNA-seq reveals alternative promoter usage as a mechanism of activating oncogenic programmes during prostate cancer progression. Nat Cell Biol. 2024 Jul;26(7):1176–86.

42. Dong Y, Liu X, Jiang B, Wei S, Xiang B, Liao R, et al. A Genome-Wide Investigation of Effects of Aberrant DNA Methylation on the Usage of Alternative Promoters in Hepatocellular Carcinoma. Front Oncol. 2022 Jan 17;11:780266.

43. Lê S, Josse J, Husson F. FactoMineR: An R Package for Multivariate Analysis. J Stat Softw. 2008 Mar 18;25:1–18.

44. Krijthe J. Rtsne: T-distributed stochastic neighbor embedding using a Barnes-Hut implementation. CRAN Contrib Packag. 2015;

45. Liberzon A, Subramanian A, Pinchback R, Thorvaldsdóttir H, Tamayo P, Mesirov JP. Molecular signatures database (MSigDB) 3.0. Bioinformatics. 2011 Jun 15;27(12):1739–40.

46. Wickham H. Introduction. In: Wickham H, editor. ggplot2: Elegant Graphics for Data Analysis [Internet]. Cham: Springer International Publishing; 2016. p. 3–10. Available from: 10.1007/978-3-319-24277-4_1

47. Thomas-Chollier M, Darbo E, Herrmann C, Defrance M, Thieffry D, van Helden J. A complete workflow for the analysis of full-size ChIP-seq (and similar) data sets using peak-motifs. Nat Protoc. 2012 Aug;7(8):1551–68.

48. Thomas-Chollier M, Herrmann C, Defrance M, Sand O, Thieffry D, van Helden J. RSAT peak-motifs: motif analysis in full-size ChIP-seq datasets. Nucleic Acids Res. 2012 Feb;40(4):e31.

49. Rauluseviciute I, Riudavets-Puig R, Blanc-Mathieu R, Castro-Mondragon JA, Ferenc K, Kumar V, et al. JASPAR 2024: 20th anniversary of the open-access database of transcription factor binding profiles. Nucleic Acids Res. 2024 Jan 5;52(D1):D174–82.

50. Heinz S, Benner C, Spann N, Bertolino E, Lin YC, Laslo P, et al. Simple combinations of lineage-determining transcription factors prime cis-regulatory elements required for macrophage and B cell identities. Mol Cell. 2010 May 28;38(4):576–89.

51. Kheradpour P, Kellis M. Systematic discovery and characterization of regulatory motifs in ENCODE TF binding experiments. Nucleic Acids Res. 2014 Mar;42(5):2976–87.

52. Kuhn M. Building Predictive Models in R Using the caret Package. J Stat Softw. 2008 Nov 10;28:1–26.

53. Hothorn T. maxstat: Maximally Selected Rank Statistics. R package version 0.7-25. 2017; Available from: https://CRAN.R-project.org/package=maxstat

54. Therneau TM. A Package for Survival Analysis in S. version 2.38. 2015; Available from: https://cran.r-project.org/package=survival

55. Therneau TM, Grambsch PM. The Cox Model. In: Therneau TM, Grambsch PM, editors. Modeling Survival Data: Extending the Cox Model. New York, NY: Springer; 2000. p. 39–77. Available from: 10.1007/978-1-4757-3294-8_3

56. Rapin N, Bagger FO, Jendholm J, Mora-Jensen H, Krogh A, Kohlmann A, et al. Comparing cancer vs normal gene expression profiles identifies new disease entities and common transcriptional programs in AML patients. Blood. 2014 Feb 6;123(6):894– 904.

57. Salgado E, Bian X, Feng A, Shim H, Liang Z. Overexpression of HDAC9 Promotes Invasion and Angiogenesis of Triple Negative Breast Cancer by regulating microRNA-206. Biochem Biophys Res Commun. 2018 Sep 5;503(2):1087–91.

58. Maccallini C, Ammazzalorso A, De Filippis B, Fantacuzzi M, Giampietro L, Amoroso R. HDAC Inhibitors for the Therapy of Triple Negative Breast Cancer. Pharmaceuticals. 2022 May 26;15(6):667.

59. Yoshida M, Kudo N, Kosono S, Ito A. Chemical and structural biology of protein lysine deacetylases. Proc Jpn Acad Ser B Phys Biol Sci. 2017 May 11;93(5):297–321.

60. Hu S, Cai J, Fang H, Chen Z, Zhang J, Cai R. RPS14 promotes the development and progression of glioma via p53 signaling pathway. Exp Cell Res. 2023 Feb 1;423(1):113451.

61. Ebert BL, Pretz J, Bosco J, Chang CY, Tamayo P, Galili N, et al. Identification of RPS14 as a 5q-syndrome gene by RNA interference screen. Nature. 2008 Jan 17;451(7176):335–9.

62. Zhou X, Hao Q, Liao J ming, Liao P, Lu H. Ribosomal Protein S14 Negatively Regulates c-Myc Activity. J Biol Chem. 2013 Jul 26;288(30):21793–801.

63. Zhou X, Hao Q, Liao J ming, Zhang Q, Lu H. Ribosomal Protein S14 Unties the MDM2-p53 Loop Upon Ribosomal Stress. Oncogene. 2013 Jan 17;32(3):388–96.

64. Schiano Lomoriello I, Giangreco G, Iavarone C, Tordonato C, Caldieri G, Serio G, et al. A self-sustaining endocytic-based loop promotes breast cancer plasticity leading to aggressiveness and pro-metastatic behavior. Nat Commun. 2020 Jun 15;11:3020.

65. Wallden B, Storhoff J, Nielsen T, Dowidar N, Schaper C, Ferree S, et al. Development and verification of the PAM50-based Prosigna breast cancer gene signature assay. BMC Med Genomics. 2015 Aug 22;8:54.

66. Prat A, Adamo B, Cheang MCU, Anders CK, Carey LA, Perou CM. Molecular Characterization of Basal-Like and Non-Basal-Like Triple-Negative Breast Cancer. The Oncologist. 2013 Feb;18(2):123–33.

67. Prat A, Pineda E, Adamo B, Galván P, Fernández A, Gaba L, et al. Clinical implications of the intrinsic molecular subtypes of breast cancer. The Breast. 2015 Nov 1;24:S26– 35.

68. Yin L, Duan JJ, Bian XW, Yu S cang. Triple-negative breast cancer molecular subtyping and treatment progress. Breast Cancer Res BCR. 2020;22:61.

69. Gong X, Du D, Deng Y, Zhou Y, Sun L, Yuan S. The structure and regulation of the E3 ubiquitin ligase HUWE1 and its biological functions in cancer. Invest New Drugs. 2020 Apr;38(2):515–24.

70. Qi L, Xu X, Qi X. The giant E3 ligase HUWE1 is linked to tumorigenesis, spermatogenesis, intellectual disability, and inflammatory diseases. Front Cell Infect Microbiol. 2022 Jul 22;12:905906.

71. Yang J, Qu T, Li Y, Ma J, Yu H. Biological role of long non-coding RNA FTX in cancer progression. Biomed Pharmacother. 2022 Sep 1;153:113446.

72. Ikegaki N, Gotoh T, Kung B, Riceberg JS, Kim DY, Zhao H, et al. De novo Identification of MIZ-1 (ZBTB17) Encoding a MYC-Interacting Zinc-Finger Protein as a New Favorable Neuroblastoma Gene. Clin Cancer Res. 2007 Oct 18;13(20):6001–9.

73. Fan Y, Kao C, Yang F, Wang F, Yin G, Wang Y, et al. Integrated Multi-Omics Analysis Model to Identify Biomarkers Associated With Prognosis of Breast Cancer. Front Oncol. 2022 Jun 10;12:899900.

74. Singh S, Kumar S, Srivastava RK, Nandi A, Thacker G, Murali H, et al. Loss of ELF5– FBXW7 stabilizes IFNGR1 to promote the growth and metastasis of triple-negative breast cancer through interferon-γ signalling. Nat Cell Biol. 2020 May;22(5):591–602.

