## Supplementary File for "Alternative Promoters Drive Transcriptomic Reprogramming and Prognostic Stratification in TNBC"

Dr. Sherry Bhalla, PhD

CSIR-Institute of Genomics and Integrative Biology (IGIB),

South Campus, Mathura Road, New Delhi, 110025, India.

### Table of content of supplementary material

*Figure S1: Analysis of promoters in the cohort using RNA-seq data.*

*Figure S2: Critical promoter switching events highlighted in TNBC-specific AAPs.*

*Figure S3: Uncovering of subtype-specific AAPs provides more detailed insights.*

*Figure S4: Activity of alternative promoters predicts patient survival.*

*Figure S5: Patient stratification based on number of events.*

*Table S1: Pathway enrichment analysis for upregulated DEGs; DRPGs and common genes.*

*Table S2: Pathway enrichment analysis for downregulated DEGs; DRPGs and common genes.*

*Table S3: List of genes having both up and downregulated DRPs.*

*Table S4: List of motifs obtained through differential motif analysis for TNBC-specific AAPs.*

*Table S5: Statistics of TNBC Type 4 and PAM50 subtypes in FUSCC and external dataset 1.*

*Table S6: List of motifs obtained through differential motif analysis for TNBC subtype-specific AAPs.*

*Table S7: List of motifs obtained through differential motif analysis for prognostic AAPs.*

*Table S8: Univariate survival analysis results for all the genes in the training and validation dataset.*

*Table S9: Univariate survival analysis results for all the promoters in the training and validation dataset.*

*Table S10: Details of the clinical features assessed through univariate survival analysis along with threshold obtained; distribution of patients in high/low categories and the obtained p-values.*

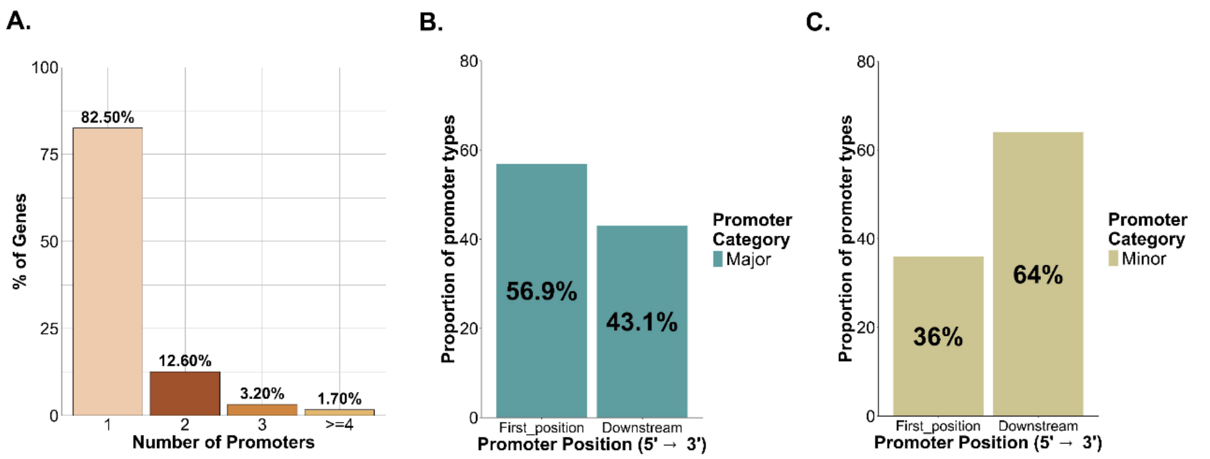


**Supplementary Fig S1. Analysis of promoters in the cohort using RNA-seq data. A.** Plot depicting the percentage of genes with a single promoter and multiple promoters. **B.** 43.1% of major promoters were positioned downstream of the first position in a gene. **C.** 64% of minor promoters were detected downstream of the first position.

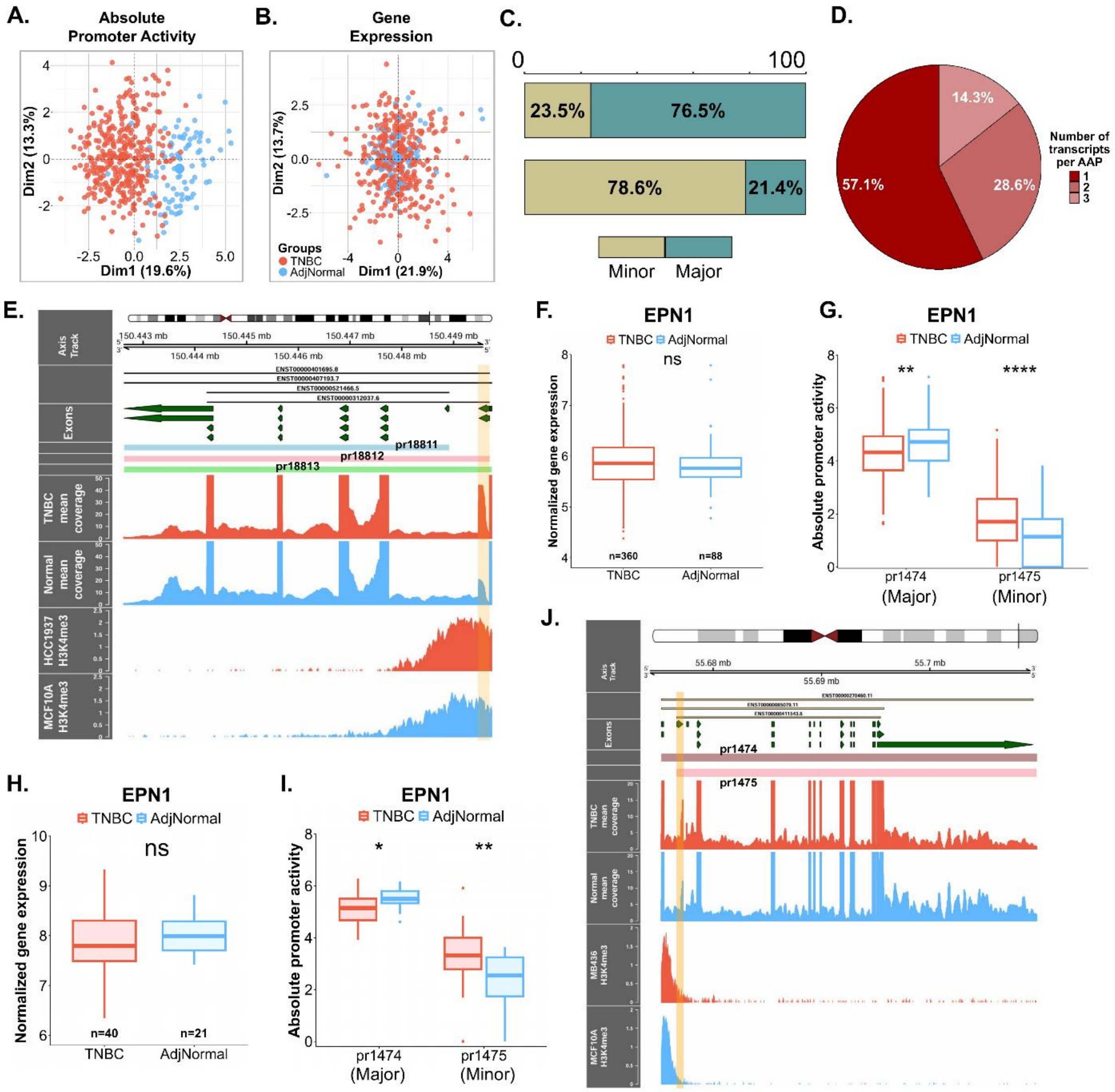


**Supplementary Fig S2. Critical promoter switching events highlighted in TNBC-specific AAPs. A.** Absolute promoter activity of TNBC AAPs is able to distinguish TNBC and normal samples. **B.** PCA plot for gene expression of TNBC-specific AAPs. **C.** A total of 23.5% minor and 76.5% major promoters were observed in FUSCC; and 78.6% of TNBC AAPs were minor promoters. **D.** Number of transcripts transcribed from the TNBC-specific AAPs. **E.** Trackplot depicting all promoters of *RPS14* gene along with their corresponding transcripts; exons; mean bam coverage and H3K4me3 peaks. **F.** *EPN1* shows non-significant gene expression in FUSCC. **G.** Minor promoter of *EPN1* shows significantly higher activity in TNBCs in FUSCC. **H & I.** Validation of gene expression **(H)** and absolute promoter activity **(I)** of *EPN1* promoters in the external dataset 1. **J.** Trackplot for *EPN1* with the mean bam coverage across TNBC and normal patients and H3K4me3 signals in cell lines.

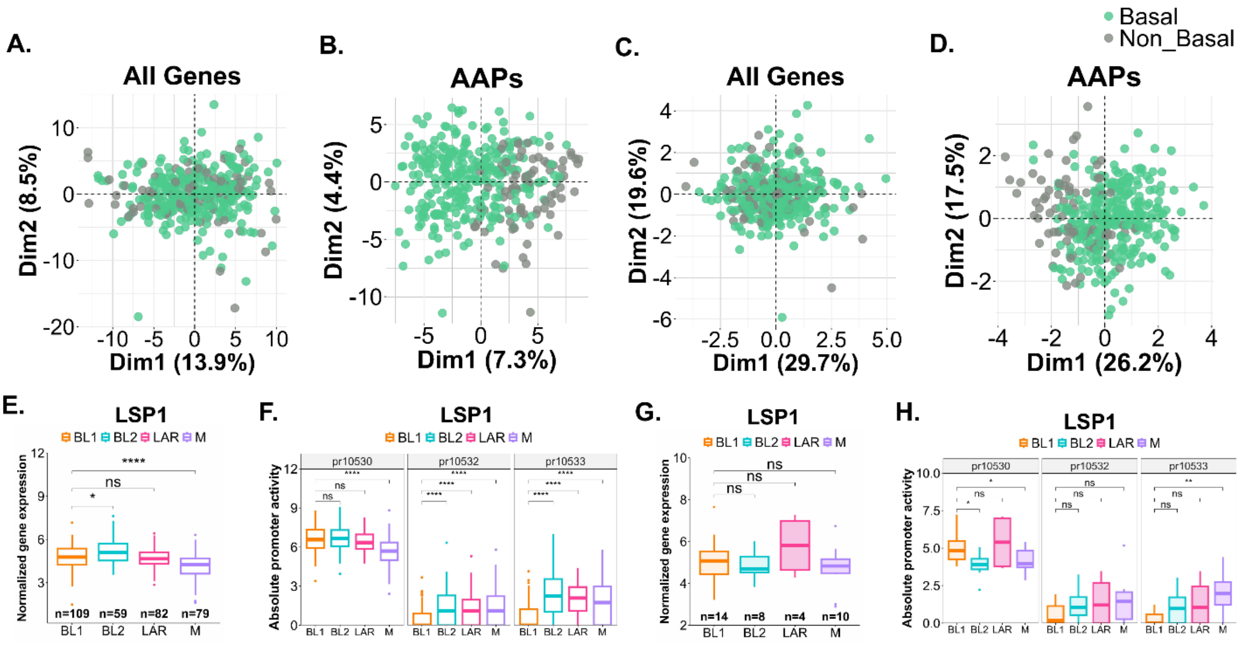


**Supplementary Fig S3. Uncovering of subtype-specific AAPs provides more detailed insights. A.** PCA plot depicting the ability of gene expression of the AAPs obtained in the FUSCC dataset in segregating the basals from non-basal samples. **B.** PCA plot for the absolute promoter activity of the FUSCC basal AAPs. **C & D.** The plots depict separation of basal and non-basal samples by gene expression **(C)** and promoter activity **(D)** of the 6 validated basal specific AAPs. **E.** Comparison of *LSP1* gene expression across all TNBC subtypes in the FUSCC dataset. **F.** *LSP1* promoter activity variation across TNBC subtypes in FUSCC. **G & H.** Boxplots showing gene expression **(G)** and promoter activity **(H)** difference across all the individual subtypes of TNBC in external dataset 1.

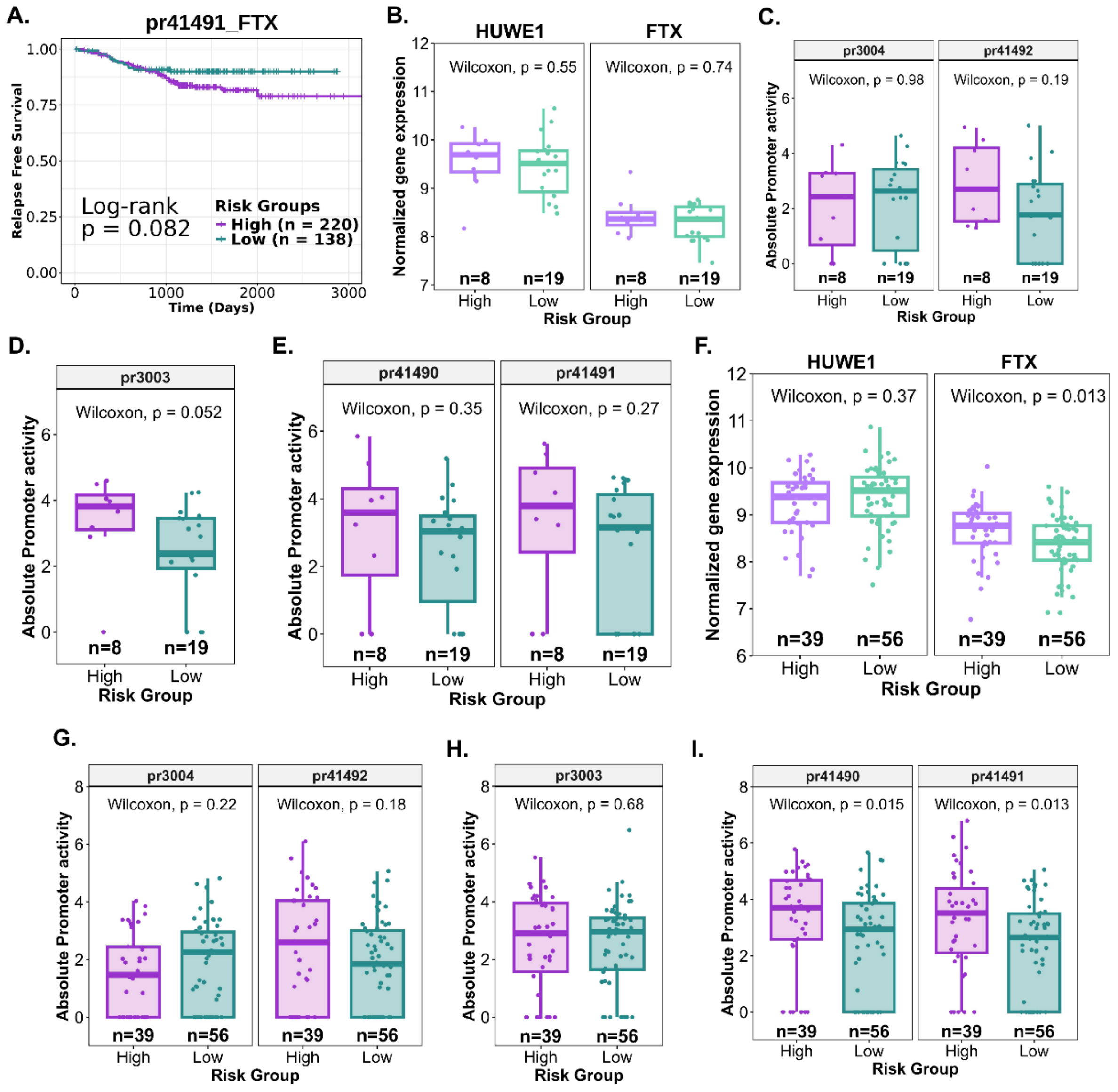


**Supplementary Fig S4. Activity of alternative promoters predicts patient survival A.** RFS based on *FTX* pr41491 promoter activity, with non-significant stratification. **B.** External validation in TNBC patients: Boxplot showing non-significant differences in *FTX* and *HUWE1* gene expression between the high-risk and low-risk patients. **C.** External validation in TNBC patients: Boxplot illustrating high pr41492 promoter activity and low pr3004 promoter activity in high-risk patients. **D & E.** Box plots depicting promoter activity for remaining *FTX* and *HUWE1* promoters. **F.** External validation of gene expression of *HUWE1* and *FTX* in all breast cancer patients. **G.** Higher activity of pr3004 in low risk and of pr41492 in high risk breast cancer patients of external dataset 3. **H & I.** External validation of other promoters of *HUWE1* and *FTX* in all breast cancer patients.

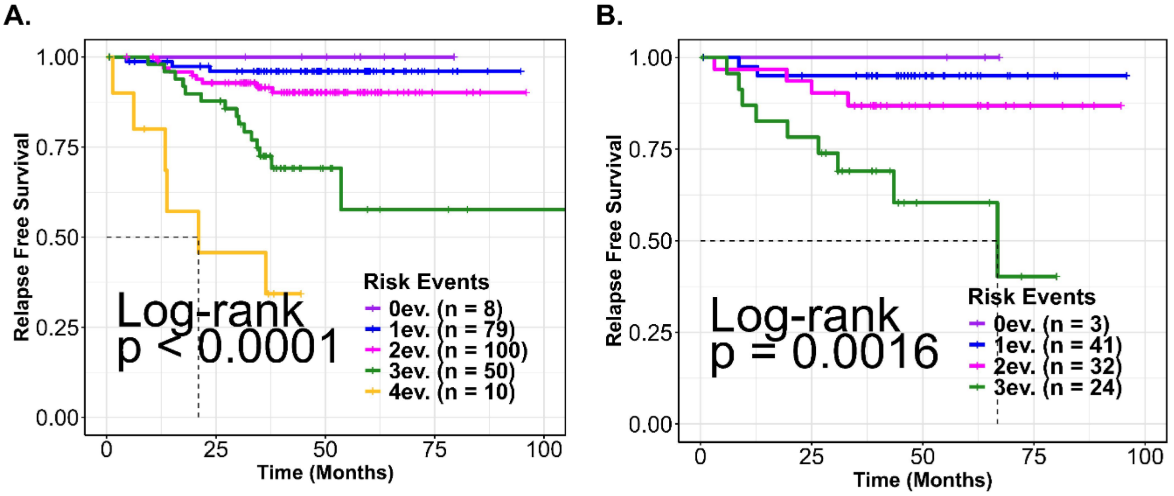


**Supplementary Fig S5. Patient stratification based on number of events.** Kaplan-Meier curves with all defined events showing RFS in the **A.** Training dataset **B.** Validation dataset.
